# Network fingerprint of stimulation-induced speech impairment in essential tremor

**DOI:** 10.1101/2020.02.20.958470

**Authors:** Jan Niklas Petry-Schmelzer, Hannah Jergas, Tabea Thies, Julia K. Steffen, Paul Reker, Haidar S. Dafsari, Doris Mücke, Gereon R. Fink, Veerle Visser-Vandewalle, Till A. Dembek, Michael T. Barbe

## Abstract

**Objective:** To gain insights into structural networks associated with stimulation-induced dysarthria (SID) and to predict stimulation-induced worsening of intelligibility in essential tremor patients with bilateral thalamic deep brain stimulation (DBS).

**Methods:** Monopolar reviews were conducted in 14 essential tremor patients. Testing included determination of SID thresholds, intelligibility ratings and a fast syllable repetition task. Volumes of tissue activated (VTAs) were calculated to identify discriminative fibers for stimulation-induced worsening of intelligibility in a structural connectome. The resulting fiber-based atlas structure was than validated in a leave-one-out design.

**Results:** Fibers determined as discriminative for stimulation-induced worsening of intelligibility were mainly connected to the ipsilateral precentral gyrus as well as to both cerebellar hemispheres and the ipsilateral brainstem. In the thalamic area, they ran laterally to the thalamus and postero-medially to the subthalamic nucleus, in close proximity, mainly antero-laterally, to fibers beneficial for tremor control as published by Al-Fatly et al. (2019). The overlap of the respective clinical stimulation setting’s VTAs with these fibers explained 62.4% (p<0.001) of the variance of stimulation-induced change in intelligibility in a leave-one out analysis.

**Interpretation:** This study demonstrates that SID in essential tremor patients is associated with both, motor cortex and cerebellar connectivity. Furthermore, the identified fiber-based atlas structure might contribute to future postoperative programming strategies to achieve optimal tremor control without speech impairment in ET patients with thalamic DBS.

## Introduction

Essential tremor is the most common adult movement disorder with an estimated prevalence of nearly 5% in populations over 65years. For medication refractory cases, deep brain stimulation (DBS) of the ventral intermediate nucleus of the thalamus (VIM) and the posterior subthalamic area (PSA) is an established, effective, and safe treatment.^1, 2^ However, side effects such as stimulation-induced dysarthria (SID), ataxia, and muscle contractions may occur.^3^ Among these, SID is one of the most common and disabling side effects, affecting up to 75% of the patients, interfering with quality of life and social functioning.^4^ Unfortunately, SID also impacts tremor control, as it often leads to suboptimal stimulation settings to avoid speech problems.

The so far hypothesized pathogenesis of SID is a modulation of cerebellothalamic fibers^5, 6^ and connections from the motor cortex, i.e., the internal capsule,^7^ leading to a functional impairment of the speech motor system and hence a deterioration of speech intelligibility. While these studies examined patients with Parkinson’s disease, previous studies by our group also showed an impairment of the speech motor system in essential tremor patients with thalamic DBS. Thalamic DBS led to an impairment of articulatory coordination patterns, probably due to a modulation of cerebellothalamic connections.^8^ Moreover, we observed articulatory imprecision and slowness affecting the production of the entire syllable cycle, most likely related to a modulation of the internal capsule.^9–11^ However, the exact pathogenesis of SID remains elusive.

Besides exploring the origin of SID, there is an urgent need to establish strategies to avoid SID while maintaining effective tremor control. Previous studies focused on the influence of lead placement and stimulation settings, and included dysarthria as a dichotomized symptom.^11, 12^ In the present study, we chose speech intelligibility as the primary outcome parameter in order to predict the deterioration of speech under DBS.

This measurement reflects real-life speech impairment and has been elaborated as a valid parameter for stimulation-induced impairment of the speech motor system.^9, 11, 13^ By combining measurements of speech intelligibility with a state-of-the-art normative structural connectome approach, we aimed to identify fibers predicting stimulation-induced worsening of intelligibility. Additionally to offering insights into the networks associated with SID, these fibers might then serve as supportive atlas structures in postoperative programming sessions to avoid the occurrence of SID while maintaining effective tremor control.

## Methods

### Patient selection and ethics

Inclusion criteria were a confirmed diagnosis of medically intractable essential tremor according to the International Parkinson and Movement Disorder Society consensus diagnostic criteria, bilateral thalamic DBS implantation (VIM or VIM/PSA) at least three months before study participation, age >18 and <80 years, and German as native language. The occurrence of postoperative SID was not an inclusion criterion, and patients were not tested for voice tremor under inactivated stimulation before study participation. The study was carried out following the Declaration of Helsinki and was approved by the local ethics committee (17-425). All patients gave written informed consent before study participation.

### Clinical assessment

First, patients were assessed in “ON stimulation” state, with their regular stimulation settings as optimized per clinical routine, and in the “OFF stimulation” state after turning the stimulation off for at least 15 minutes. Second, a hemisphere-wise monopolar review was conducted with stimulation of the contralateral hemisphere turned off. When leads with eight contact heights had been implanted (Vercise™ leads, Boston Scientific), only every second contact, beginning at the tip, was included in the testing to increase spatial distribution and to diminish the examination time needed. All other leads (Cartesia™ leads, Boston Scientific, Medtronic 3387/3389 leads, Medtronic) were tested at each of the four contact heights, with circular stimulation mode for directional contacts. Contacts were tested in randomized order to maintain blinding of the examiner and the patient to the order of the tested contacts. The frequency was set to 130 Hz and a pulse width of 60 μs was chosen for every patient. Before testing, impedances were measured to ensure the integrity of the DBS system and constant current mode was set if applicable.

Monopolar reviews were performed by increasing the stimulation amplitude in steps of 0.5 mA until (i) a maximum of 10mA, (ii) the occurrence of intolerable side effects, or (iii) the onset of SID. The onset of SID was determined by a trained examiner (JNPS), asking the patient to enumerate the names of the months repeatedly at each amplitude step. Furthermore, contralateral muscle contractions and ataxia, defined as contralateral dysmetria during the finger-to-nose test, were documented.

The following clinical assessments were conducted in the (i) “ON stimulation” state, (ii) “OFF stimulation” state, and (iii) at each contact’s maximum stimulation amplitude as defined during the monopolar review.

#### Tremor assessment

In the “ON stimulation” and “OFF stimulation” state, tremor severity was measured based on the Tremor Rating Scale (TRS). To shorten the duration of testing, tremor severity during the monopolar review was determined by the rater as postural tremor control of the contralateral arm, rated on a visual analogue scale (VAS) ranging from 0 cm (“no tremor”) to 10 cm (“most severe tremor”).

#### Speech assessment

In each condition, patients were asked to enumerate the months and rate their “ability to speak” on a VAS ranging from 0 cm (“normal”) to 10 cm (‘worst”). The examiner was kept blinded to the patient’s VAS rating throughout the testing. Then acoustic recordings were conducted in a sound-attenuated booth. Speech patterns are highly sensitive to contextual variability on the segmental and prosodic level. To obtain a controlled corpus that was suited for our quantitative analysis, and concurrently time-saving enough to be used in a test design of multiple stimulation settings, we reduced our analysis to a fast syllable repetition task (oral diadochokinesis, DDK) and a short reading passage. DDK is a standard task in which speakers were instructed to produce consonant-vowel-sequences as clearly, quickly and often as possible in one single breath. The examiner demonstrated the task before the beginning of the first recording. Syllables with velar stop consonants requiring complex tongue body movements, /kakaka/, were used, as it has been shown that they reliably reflect stimulation-induced impairments of glottal and oral speech motor control in essential tremor patients treated with DBS.^9, 11^ For natural sentence production patients were instructed to read a short passage, the German standard text (“Nordwind und Sonne”/ “Northwind and Sun”) at a comfortable speech rate.^11, 13^

#### Intelligibility ratings and DDK analysis

To investigate speech intelligibility perceived by naïve listeners (all German native speakers), we extracted the sentence with the fewest reading errors from the natural sentence production task as auditive stimuli. We chose the sentence with the highest reading fluency to avoid effects of speech errors on the segmental and prosodic level on the intelligibility ratings and to increase the comparability of test sentences between the different stimulation settings.^14–16^ These were normalized to the same overall intensity level to minimize an effect of stimulus loudness on perception results. Eleven naïve listeners rated the stimuli in a randomized order across the conditions but matched for patients to ensure blinding of the listeners to the stimulation condition. Each stimulus was evaluated on a VAS reaching from 1 point (“very bad intelligibility”) to 101 points (“very good intelligibility”). For further analysis intelligibility rating per stimuli was calculated as the mean of all ratings across listeners.

The DDK labeling was carried out in a blinded manner by an experienced phonetician (TT), as described previously.^11^ In brief, syllable durations for each /kakaka/ sequence were identified from the start of the consonantal closure to the end of the vocalic opening with respect to the energy in the higher formant structure in the acoustic waveform. We used syllable duration as the most robust acoustic parameter described in the literature when objectively capturing the degree of motor speech impairment in DDKs.^13, 17^ Prolonged syllable durations reflect an overall slowing down of the speech motor system and can be directly related to motor speech impairment.^18^ There is a debate in the literature on whether DDKs are comparable to natural sentence production.^19^ However, in a study on effects of thalamic deep brain stimulation on speech production in ET patients, Hermes et al. have shown, that slowing down of the speech motor system can be attested for both, DDK tasks and natural sentence production.^20^ Furthermore, syllable duration has previously been shown to have a significant effect on intelligibility ratings and to be worsened by thalamic DBS in essential tremor patients.^11, 13^

### Localization of DBS leads and volume of tissue activated estimation

DBS leads were localized with the Lead-DBS toolbox (www.lead-dbs.org).^21^ The detailed processing pipeline has been described elsewhere.^22^ In brief, postoperative CT images (IQon Spectral CT, iCT 256, Brilliance 256, Philips Healthcare) were linearly co-registered to preoperative MRI (3T Ingenia, Achieva, 1.5T Ingenia, Philips Healthcare) using advanced normalization tools (ANTs, http://stnava.github.io/ANTs/, N = 13)^23^ or BRAINSFIT (https://github.com/BRAINSia/BRAINSTools, N = 1)^24^ if ANTs did not provide sufficient results after visual inspection.^25^ Then images were nonlinearly normalized into standard space (ICBM 2009b NLIN, Asym) using ANTs and the “effective (low variance)” strategy.^26^ DBS leads were automatically pre-reconstructed with the PaCER-algorithm,^27^ manually refined, and corrected for postoperative brain shift as implemented in Lead-DBS. The orientation of directional leads was determined by the DiODE algorithm.^28^

For each stimulation setting, volumes of tissue activated (VTA) were calculated in the patient’s native space and then transformed into standard space based on the individual nonlinear normalization. A well-established finite element method, introduced by Horn et al.^22^, was employed to estimate the spread of the electric field for homogenous tissue with a conductivity of σ = 0.1 S/m.^29^ The VTA was thresholded at the electrical field isolevel of 0.19 V/mm^29^ to reflect clinical stimulation results of Mädler and Coenen et al.^30^, adapted depending on the respective pulse width.^31^ Finally, right-sided VTAs were nonlinearly flipped to the left hemisphere using Lead-DBS for further analysis.^32^

### Determination of discriminative fibers

Determination of discriminative fibers for stimulation-induced worsening of intelligibility followed an approach introduced by Baldermann et al.^33^ based on a normative structural connectome.^34^ This structural connectome has previously been used to successfully predict tremor outcomes after thalamic DBS in essential tremor patients^32^ and is based on diffusion data collected in 20 subjects using a single-shot spin-echo planar imaging sequence (repetition time = 10.000ms, echo time = 94ms, 2×2×2mm^3^, 69 slices). Global fiber-tracking was performed using Gibb’s tracking method,^35^ and the resulting fibers were warped into standard space.^34^ In the present study, each of these fibers’ discriminative value was tested by associating the fiber’s connectivity to VTAs across patients with the respective change in intelligibility. Specifically, a linear mixed effect model was set up for each fiber connected to at least 20 VTAs, to test for differences between changes in speech intelligibility of connected and unconnected VTAs. The grouping variable (unconnected/connected) was included as a fixed-effect and the respective lead and patient were included as random-effects, to reflect that two leads per patient and a total of 28 leads were tested. Change scores were calculated as percentage change from “OFF stimulation” state ((intelligibility_endpoint_)–intelligibility_”OFFstimulation”_)/intelligibility_”OFFstimulation”_) and divided by the respective amplitude as proposed by Dembek et al.^31^, as changes in intelligibility in “ON stimulation state” were highly correlated with stimulation amplitude (rho = −0.66, p = 0.01). When connected VTAs led to worsening of speech intelligibility in comparison to unconnected VTAs with a p-value below 0.05, a fiber was determined as discriminative. Importantly, “ON stimulation” state VTAs were not included in the determination of discriminative fibers.

### Validation of discriminative fibers

Discriminative fibers were validated in a leave-one-out design.^22, 32, 33^ Briefly, discriminative fibers were recalculated for each patient based on the clinical assessments of all other patients. Then the respective number of discriminative fibers stimulated in the left-out patient’s “ON stimulation” state was summed up to conduct a linear regression analysis. This was done to investigate the influence of the amount of discriminative fibers stimulated on the percentage change of intelligibility in the “ON stimulation” state. An additional linear regression analysis was conducted to investigate the influence of the amount of discriminative “speech”-fibers stimulated on postoperative tremor control, defined as the change in TRS hemiscores in “ON stimulation” compared to “OFF stimulation”.

## Results

### Patient characteristics

Fourteen patients diagnosed with essential tremor (four female) aged 65.4 (±13) years were prospectively recruited, resulting in 28 leads and 112 contacts investigated. Lead types and clinical stimulation settings are described in Table 1. All patients were right-handed and bilaterally implanted either targeting the VIM (N = 2) or the VIM/PSA (N = 12) with a mean time of 29.2 (±24) months since implantation.

**Table 1.** Clinical stimulation settings.

| Patient | Lead type | Clinical stimulation settings |  |
| --- | --- | --- | --- |
|  |  | Left hemisphere | Right hemisphere |
| 1 | Cartesia™,<br>Boston Scientific | C+, 2- 21%, 3- 5%, 4- 29%, 5- 17%, 6- 5%, 7- 23%, 60µs, 149Hz, 2.3mA | C+, 10- 17%, 11- 26%, 12- 27%, 13- 10%, 14- 10%, 15- 10%, 60µs, 149Hz, 2.3mA |
| 2 | Cartesia™,<br>Boston Scientific | 5+ 34%, 6+ 33%, 7+ 33%, 1- 90%, 2- 5%, 4- 5%, 40µs, 204Hz, 5.0mA | 13+ 34%, 14+ 33%, 15+ 33%, 9- 80%, 11- 10%, 12- 10%, 40µs, 204Hz, 4.7mA |
| 3 | Medtronic 3387,<br>Medtronic | C+, 1- 100%, 60µs, 180Hz, 3.0mA | C+, 9- 100%, 60µs, 180Hz, 1.9mA |
| 4 | Cartesia™,<br>Boston Scientific | C+, 2- 100%, 60µs, 130Hz, 3.6mA | C+, 10- 29%, 11- 43%, 12- 28%, 60µs, 130Hz, 1.9mA |
| 5 | Cartesia™,<br>Boston Scientific | C+, 2- 34%, 3- 33%, 4- 33%, 60µs, 130Hz, 2.1mA | C+, 10- 34%, 11- 33%, 12- 33%, 60µs, 130Hz, 2.1mA |
| 6 | Vercise™,<br>Boston Scientific | C+, 3- 100%, 60µs, 130Hz, 3.2mA | C+, 11- 100%, 60µs, 174Hz, 2.3mA |
| 7 | Medtronic 3389,<br>Medtronic | C+, 2- 100%, 60µs, 130Hz, 3.7mA | C+, 10- 100%, 60µs, 130Hz, 2.0mA |
| 8 | Vercise™,<br>Boston Scientific | C+, 3- 100%, 60µs, 174 Hz, 1.3mA | C+, 11- 100%, 60µs, 174Hz, 0.7mA |
| 9 | Vercise™,<br>Boston Scientific | C+, 3- 100%, 60µs, 130Hz, 3.8mA | C+, 11-, 100%, 60µs, 130Hz, 2.9mA |
| 10 | Medtronic 3389,<br>Medtronic | Vim 1: C+, 0-, 2.0V, 60µs, 125Hz<br>Vim 2: C+, 1-, 2.5V, 60µs, 125Hz | Vim 1: C+, 8-, 2.0V, 60µs, 125Hz<br>Vim 2: C+, 9-, 2.7V, 60µs, 125 Hz |
| 11 | Vercise™,<br>Boston Scientific | 5+ 100%, 2- 70%, 3- 30%, 60µs, 174Hz, 2.1mA | 13+ 100%, 10- 70%, 11- 30%, 60µs, 174Hz, 2.1mA |
| 12 | Cartesia™,<br>Boston Scientific | C+, 1- 100%, 50µs, 185Hz, 1.4mA | C+, 13- 35%, 14- 35%, 15- 30%, 60µs, 185Hz, 1,2mA |
| 13 | Cartesia™,<br>Boston Scientific | C+, 2- 34%, 3- 33%, 4- 33%, 60µs, 185Hz, 1.2mA | C+, 10- 34%, 11- 33%, 12- 33%, 60µs, 185Hz, 1.6mA |
| 14 | Cartesia™,<br>Boston Scientific | C+, 1- 100%, 60µs, 130 Hz, 1.2mA | C+, 9- 100%, 60µs, 130Hz, 1.2mA |

### Comparison of “OFF stimulation” and “ON stimulation” state

The mean rating of intelligibility did not differ significantly between “OFF stimulation” and “ON stimulation” state on the group level (mean OFF: 70.9 ±14.3, mean ON: 67.5 ±17.1, p = 0.763, Fig.1 A1). However, on the individual level, a formal worsening of intelligibility rating was observed in half of the patients. Regarding impairment of the speech motor system, syllable durations increased significantly during “ON stimulation” (mean OFF: 243ms ±58, mean ON: 275ms ±72, p = 0.011, Fig.1 B1). Regarding tremor control, TRS total scores improved significantly with clinical stimulation settings (mean OFF: 35.2 ±18.4, mean ON: 13.0 ±10.4, p<0.001, Fig.1 C1), reflecting a tremor suppression of over 60%.

**Figure 1.**
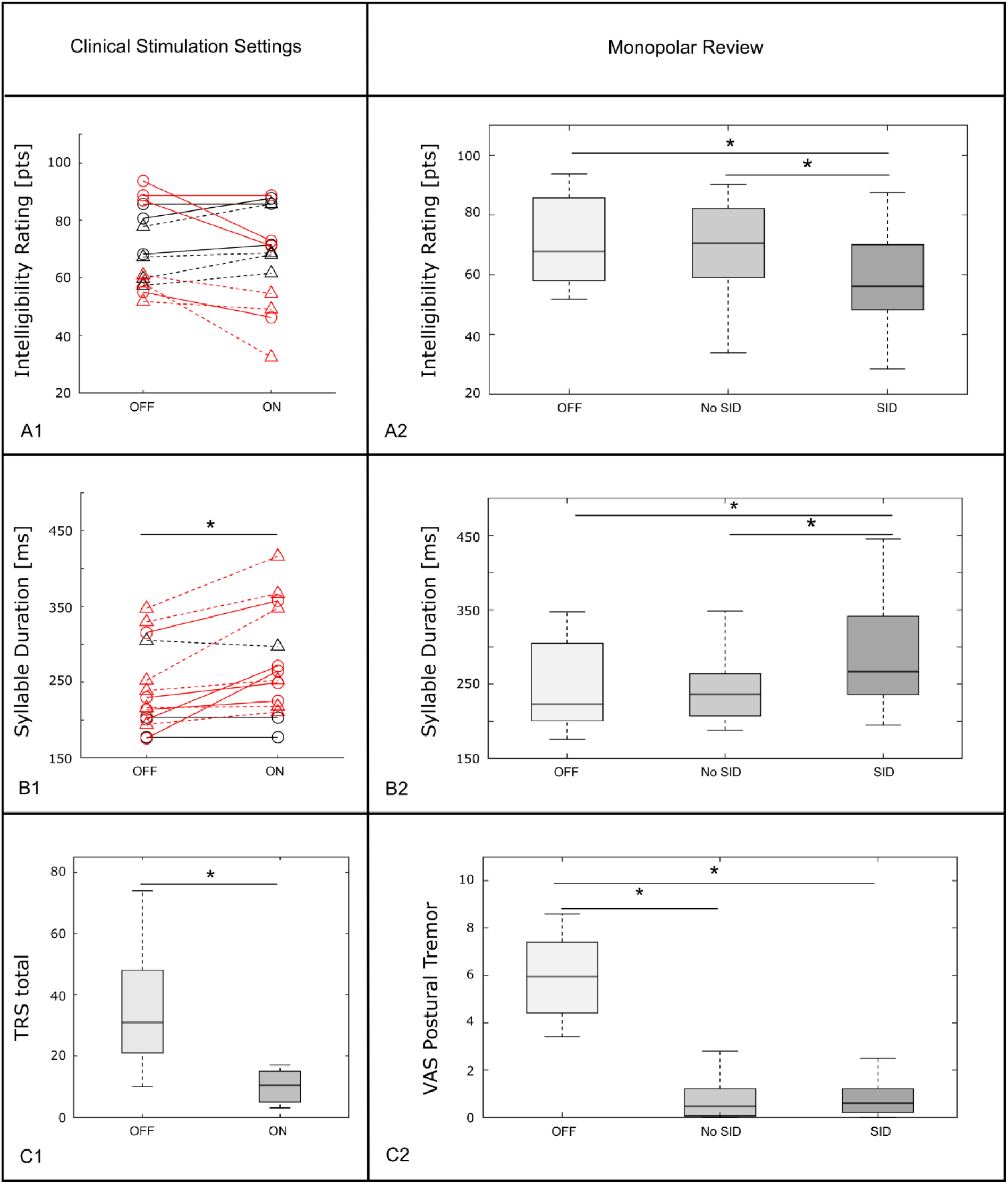
Clinical assessments. Comparison of “OFF stimulation” and “ON stimulation” states (column one) and comparison of “OFF stimulation” state, stimulation settings not causing stimulation-induced dysarthria (SID) and stimulation settings causing SID during monopolar review (column two) for intelligibility rating (A), syllable duration (B) and tremor ratings (C). Black stars indicate significant differences (p<0.05) between the settings. Triangles in column one indicate the appearance of voice tremor in the “OFF stimulation” state and individual formal worsening is marked in red. **Abbreviations: SID** = stimulation-induced dysarthria, **TRS** = tremor rating scale; **VAS** = visual analog scale

### Monopolar review outcomes

Figure 2 illustrates the anatomical position of investigated contacts and the stimulated region covering the VIM and the PSA. A total of 111 stimulation settings were included for further analysis (see Fig. 3). When comparing the “OFF stimulation” state to stimulation settings not causing SID and stimulation settings causing SID, Kruskal-Wallis test revealed differences in intelligibility ratings (X^2^(2) = 14.1, p<0.001, Fig.1 A2). Post-hoc Wilcoxon rank-sum tests showed significant differences in “OFF” versus “SID” (mean “SID”: 58.2 ±15.6, p = 0.007) and “no SID” versus “SID” (mean “no SID”: 68.5 ±14.7, p<0.001). Additionally, for syllable durations Kruskal-Wallis test (X^2^(2) = 15.6, p<0.001) and post-hoc Wilcoxon rank-sum tests also revealed significant differences for “OFF” versus “SID” (mean “SID”: 293ms ±72, p = 0.011) and “no SID” versus “SID” (mean “no SID”: 247ms ±51, p<0.001). When assessing the ability to speak, using the VAS, there was a worsening from “no SID” to “SID” (mean “no SID”: 2.0 ±2.0, mean “SID”: 3.9 ±2.0, p<0.001). When assessing postural tremor control, using the VAS, data suggested significant improvement from “OFF” to “no SID” and “OFF” to “SID” (mean “OFF”: 5.9 ±1.6, mean “no SID”: 1.0 ±1.3, p<0.001; mean “SID”: 1.0 ±1.1, p<0.001). Intelligibility ratings significantly correlated to the patients’ VAS rating of their ability to speak (rho = −0.45, p<0.001), and syllable durations (rho = −0.66, p<0.001). As shown in Figure 3, onset of SID was associated with both, muscle contractions and/or ataxia, but also occurred without any additional symptoms.

**Figure 2.**
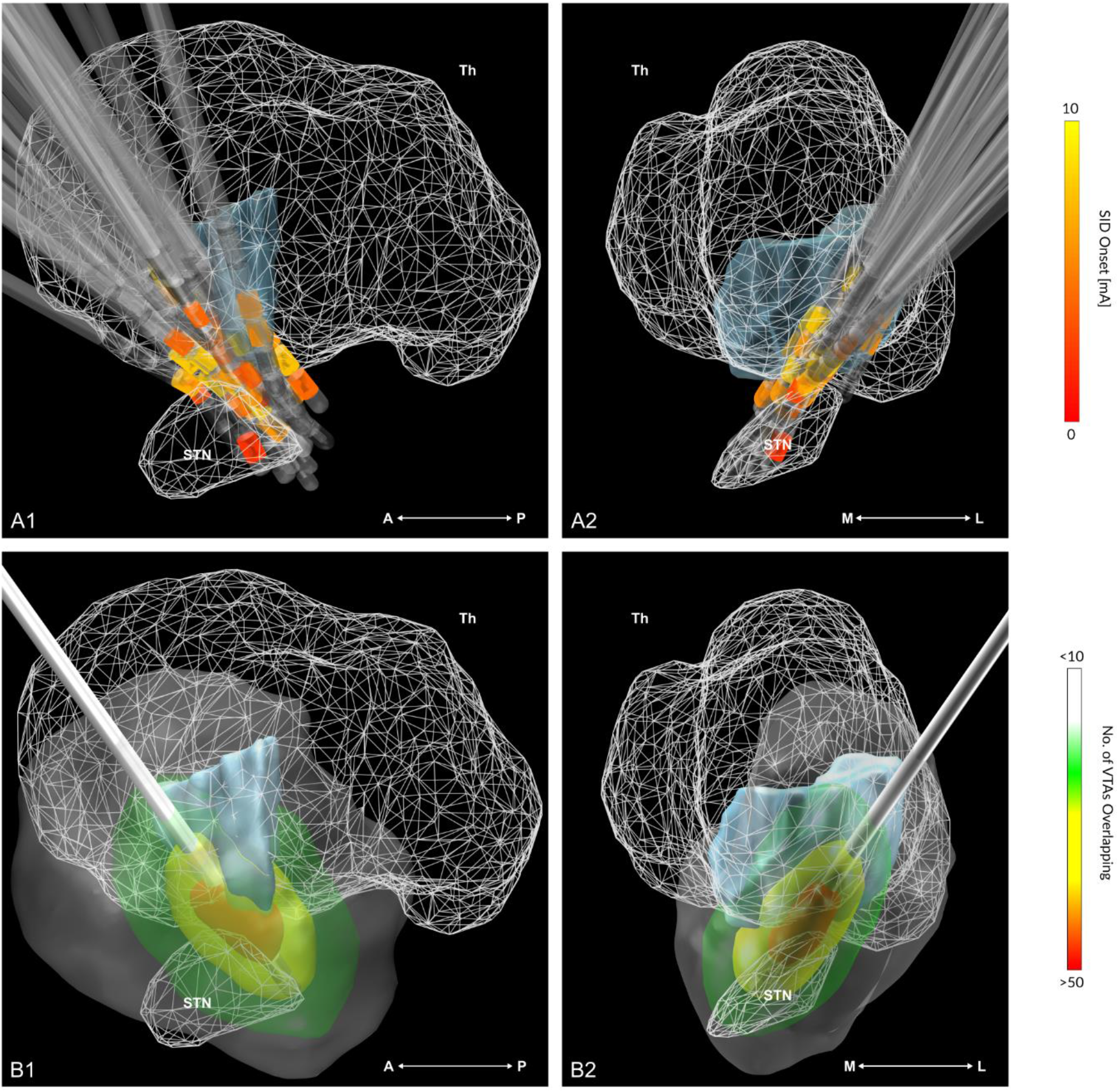
Contact positions and investigated anatomical area. (A) shows the positions of left-pooled contacts included in the analysis in standard space (ICBM 2009b NLIN, Asym) in lateral (A1) and anterior view (A2) together with the thalamus, the VIM (blue), and the STN. Contacts at which stimulation-induced dysarthria could be provoked are color-coded by the amplitude at the onset of dysarthria. (B) illustrates the investigated anatomical area together with the mean lead position in lateral (B1) and anterior view (B2), together with the thalamus, the VIM (blue), and the STN. Anatomical structures are taken from the DISTAL atlas, as included in LEAD-DBS.^21, 48^ **Abbreviations: A** = anterior, **L** = lateral, **M** = medial, **P** = posterior, **STN** = nucleus subthalamicus, **Th** = thalamus, **VIM** = ventral intermediate nucleus, **VTA** = volume of tissue activated

**Figure 3.**
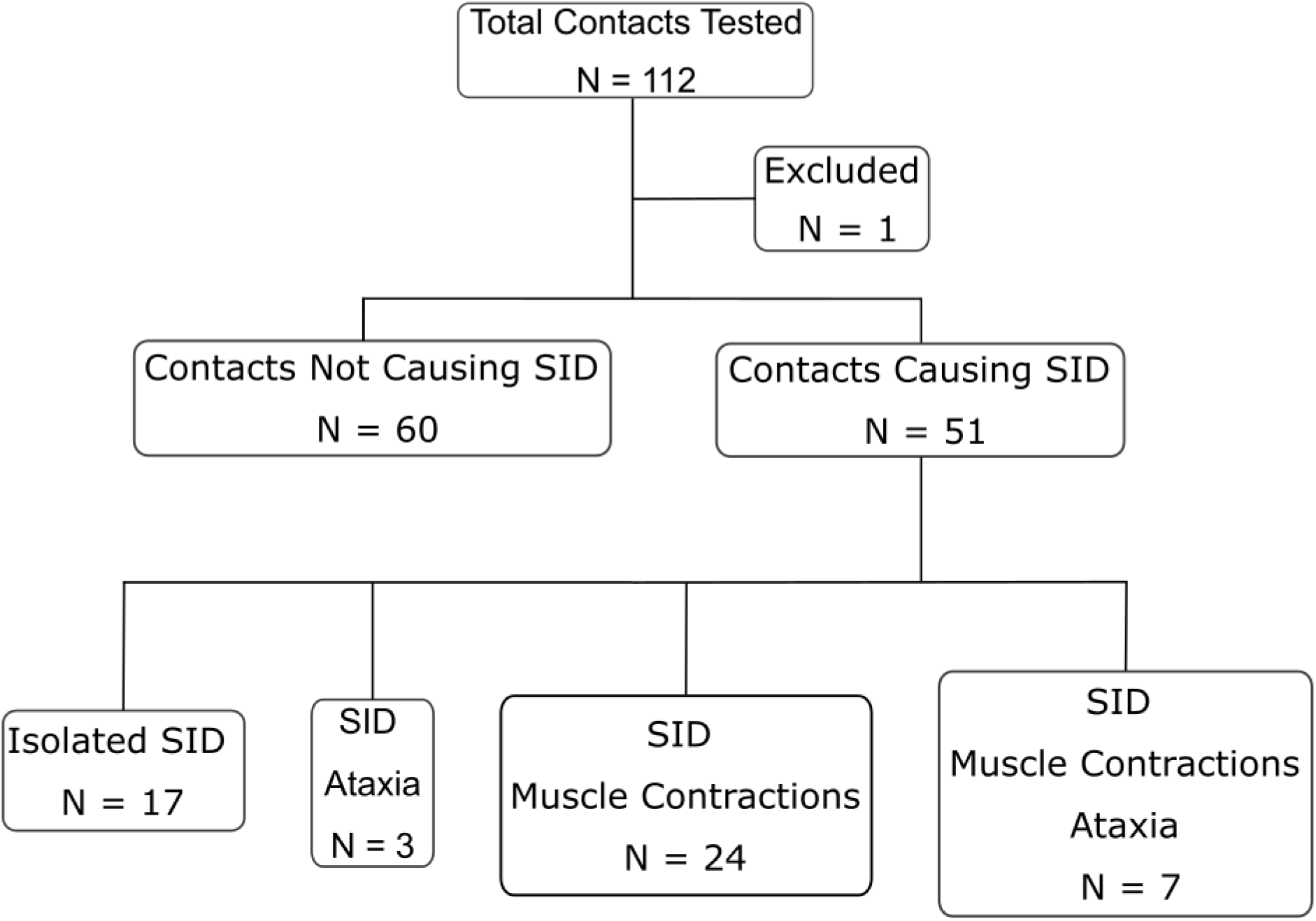
Clinical symptoms associated with stimulation-induced dysarthria. Clinical symptoms of motor cortex (muscle contractions) or cerebellar cortex (ataxia) involvement at the onset of stimulation-induced dysarthria (SID). One contact was excluded as the patient experienced side effects severely interfering with further testing. **Abbreviations: SID** = stimulation-induced dysarthria

### Discriminative fibers for worsening of intelligibility

Fibers determined as discriminative for stimulation-induced worsening of intelligibility were mainly connected to the ipsilateral precentral gyrus as well as to both cerebellar hemispheres and the ipsilateral brainstem (see Fig. 4). In the target area, they ran laterally to the thalamus and postero-medially to the subthalamic nucleus, in close proximity, mainly antero-laterally, to fibers beneficial for tremor control as published by Al-Fatly et al. ^32^. Some of them even overlapped (see Fig. 5). Validation of the discriminative fibers in a leave-one-out design revealed that the overlap of the VTAs with the discriminative fibers identified in the present study could explain 62.4% of the variance in individual intelligibility outcome with clinical stimulation settings as measured in the respective “ON stimulation” state (R^2^ = 0.624, p < 0.001, Fig. 4). Overlap of the VTAs with these discriminative “speech”-fibers was not associated with postoperative tremor control (R^2^ = 0.04, p = 0.289).

**Figure 4.**
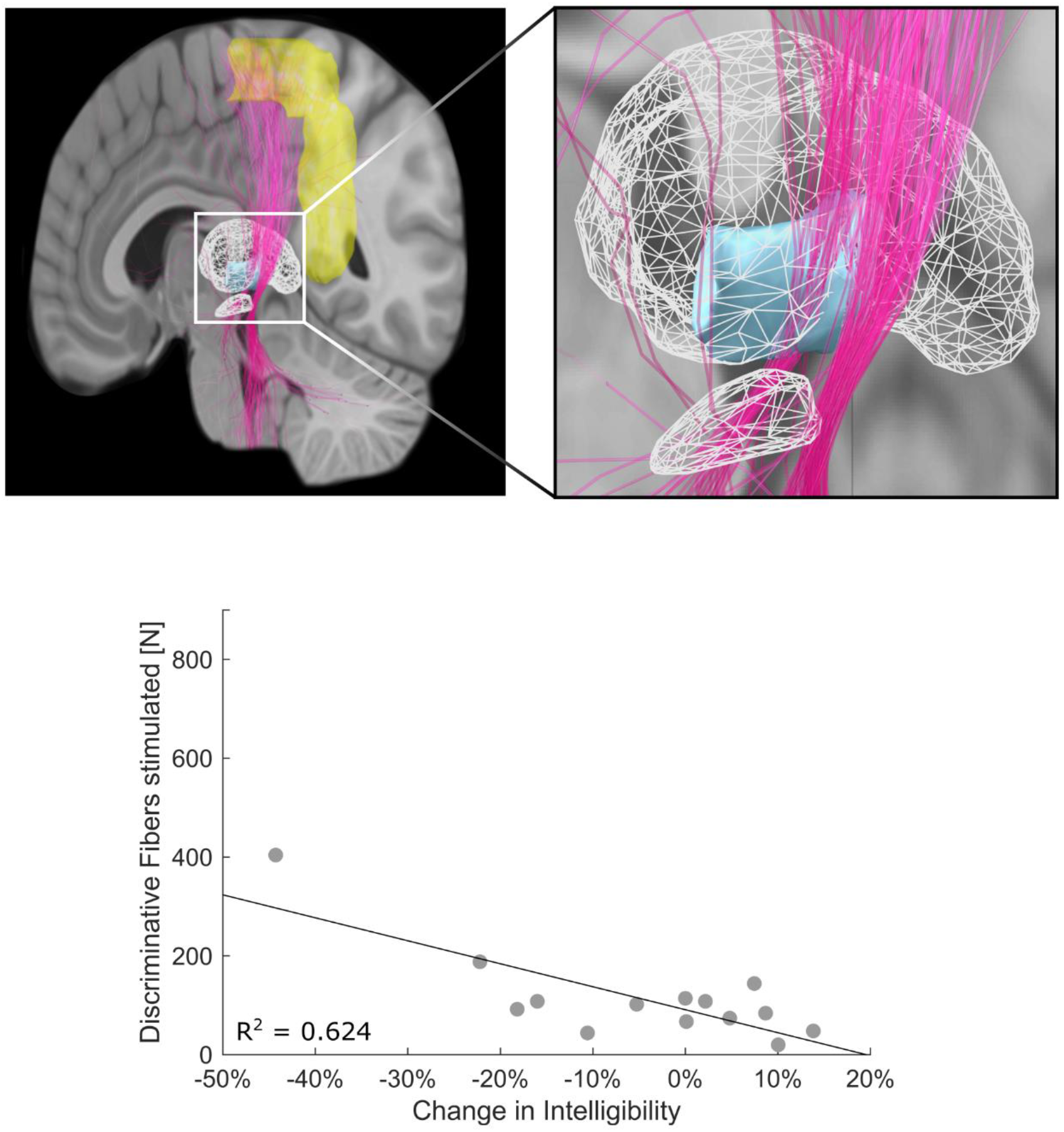
Discriminative fibers for speech intelligibility. The course of discriminative fibers for stimulation-induced changes in intelligibility (pink) as identified in a high-definition structural connectome.^34^ Fibers are shown in the antero-lateral view together with the thalamus, the VIM (blue), the precentral gyrus (yellow), and the STN. Discriminative fibers were validated in a leave-one-out design, explaining 62.4% of the variance in change of intelligibility during “ON stimulation” (p<0.001). Anatomical structures are taken from the DISTAL atlas, as included in LEAD-DBS.^21, 48^ **Abbreviations: STN** = nucleus subthalamicus, **VIM** = ventral intermediate nucleus

**Figure 5.**
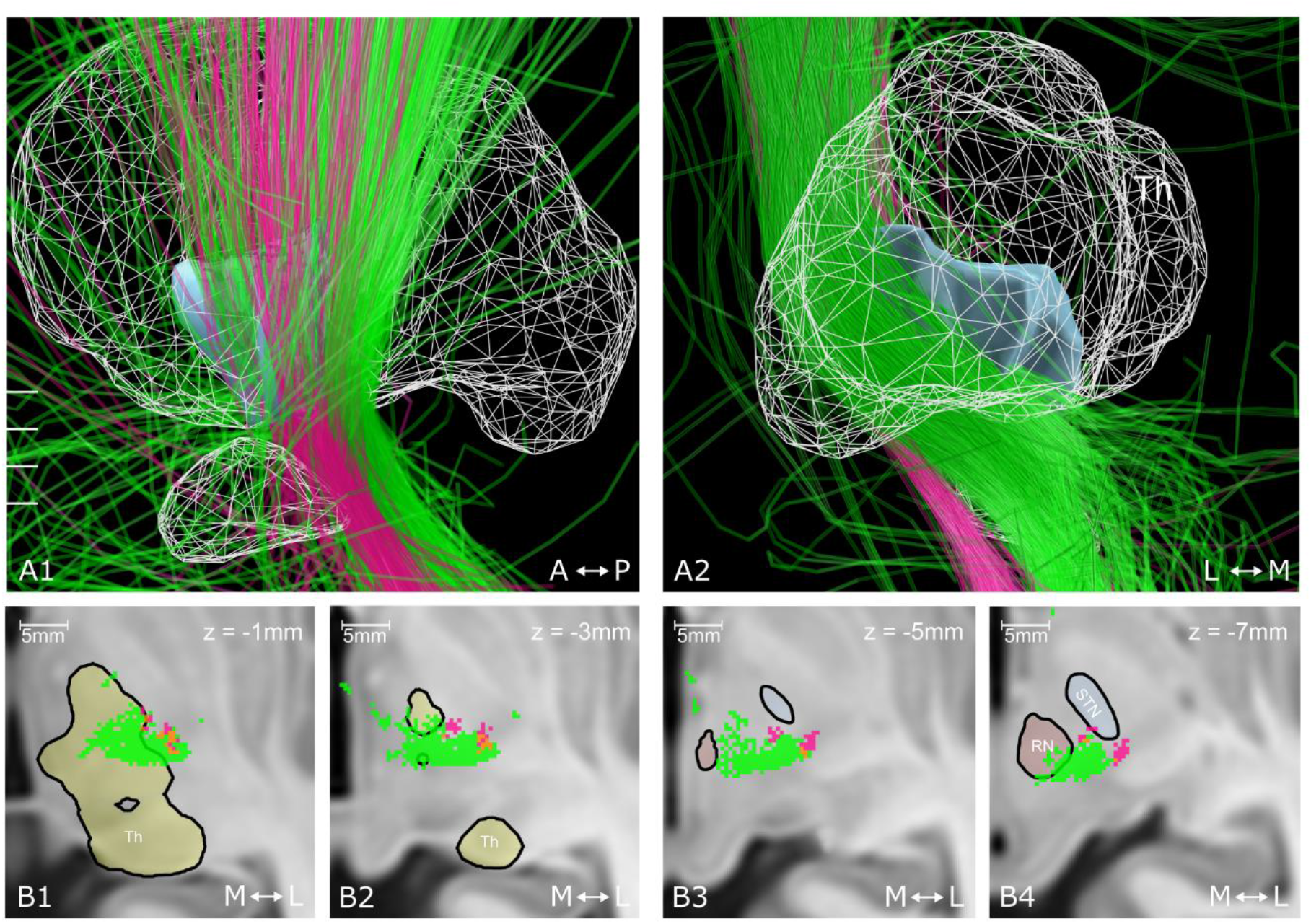
Comparison of discriminative fibers for changes of intelligibility and tremor control. The course of discriminative fibers for changes of intelligibility (pink) and postoperative tremor control (green), as identified by Al-Fatly et al.,^32^ in lateral (A1) and posterior view (A2), together with the thalamus (yellow), the VIM (blue), and the STN (light grey). (B) illustrates the course of the respective fibers in axial view with overlapping areas marked in orange. Slice positions are indicated in A1. For increased clarity only voxels with at least six neighbored voxels containing the respective fiber tracts are shown. Anatomical structures are taken from the DISTAL atlas, as included in LEAD-DBS.^21, 48^ **Abbreviations: A** = anterior, **L** = lateral, **M** = medial, **P** = posterior, **RN** = red nucleus, **STN =** nucleus subthalamicus, **Th** = thalamus, **VIM** = ventral intermediate nucleus

## Discussion

This study demonstrates that stimulation-induced speech impairment in thalamic DBS for essential tremor patients is associated with both, motor cortex and cerebellar connectivity. Furthermore, the identified discriminative fibers were predictive for worsening of intelligibility with clinical stimulation settings in our cohort.

### Discriminative fibers to predict worsening of intelligibility

Discriminative fibers for stimulation-induced worsening of intelligibility were identified to run laterally to the motor thalamus and postero-medially to the subthalamic nucleus. These fibers were either nearby, mainly antero-laterally, or even overlapping with fibers beneficial for tremor control as published by Al-Fatly et al..^32^ The overlap of the clinical stimulation settings with these fibers was predictive of 62.4% of the variance of stimulation-induced changes of intelligibility, as revealed in a leave-one-out design (see Fig. 4 and 5).

Intelligibility was chosen as a parameter representing the degree of SID in essential tremor patients since it indicates a deterioration of the speech motor system and represents the patient-rated “ability to speak”, and is therefore of high clinical relevance in daily living.^11^ In contrast to previous studies, no worsening of intelligibility rating was observed with bilateral DBS on the group-level when comparing “On stimulation” and “Off stimulation”.^11, 13^ This finding might be due to the fact that some patients in the present cohort had a voice tremor during “OFF stimulation” which has been reported to improve with DBS,^36^ and thereby might have led to an improvement of intelligibility beyond its worsening due to an affection of the glottal speech motor system (see Fig. 1). Nevertheless, on the individual level, 50% of the patients experienced a worsening of intelligibility with clinical stimulation settings and there was a significant increase in syllable duration under DBS, indicating a systematic affection of the speech motor system in terms of oral control in our cohort. Of note, the identified fibers seem to be specific for changes in intelligibility as they were not predictive for tremor control.

### Pathophysiological considerations

Following Fuertinger et al.^37^ and Guenther et al.^38^ the functional connectome of real-life speech production in right-handed healthy subjects is constituted by a core hub network, centered on the left laryngeal and orofacial regions of the primary motor cortex and its main input regions in the surrounding premotor, somatosensory and parietal cortices. This core sensorimotor hub network is then widely connected to other brain regions such as the auditory cortex as an auditory feedback control subsystem, the parietal cortex for phonological and semantic processing, or the cerebellum for modulation of vocal motor timing and sequence processing.^37–39^

As expected and hypothesized in previous studies,^5–9, 11^ and in line with the described connectome of real-life speech production the identified discriminative fibers mainly connected the motor cortex to the brainstem and both cerebellar hemispheres. The association between stimulation-induced speech impairment and these cortical areas is underlined by the fact that the onset of SID was in most of the cases associated with clinical symptoms of motor cortex modulation, i.e., muscle contractions, or cerebellar cortex involvement, i.e., ataxia (see Fig. 3). The course of these fibers also fits the finding of previous studies, reporting that a more lateral contact position^11^ and contacts located outside the motor thalamus might be associated with SID in essential tremor patients.^12^ Additionally, damage to the cerebellum, especially the paravermal region, has previously been reported to result in a slurred and poorly coordinated speech,^40^ as also observed in essential tremor patients with deep brain stimulation.^8, 10^ Although indicating stimulation-induced speech impairment by cerebellar involvement, the present finding cannot be predictive of later cerebellar syndrome induction, as discussed in essential tremor patients with thalamic DBS.^41^ Previous studies investigating changes in intelligibility after DBS of the subthalamic nucleus in Parkinson’s disease provided evidence that both, modulation of motor fibers running in the internal capsule and a modulation of cerebello-thalamic pathways, i.e., the dentatorubrothalamic tract, can induce stimulation-induced speech worsening.^6, 7^ Interestingly, the course of the discriminative fibers is in line with a study by Aström et al., reporting a posteromedial contact position in relation to the subthalamic nucleus to be associated with impairment of speech intelligibility in patients with Parkinson’s disease treated with DBS of the subthalamic nucleus.^5^ Especially in the posterior subthalamic area, the identified discriminative fibers were running quite strictly separated anterior and lateral to the fibers associated with tremor control (see Fig. 5). However, this study does not provide specific connectivity patterns to the cerebellar cortex or the motor cortex, differentiating between fibers associated with tremor control or fibers predicting stimulation-induced speech impairment. Taking together the results of the present study and of previous studies providing evidence for a combination of spastic and atactic changes in the speech motor system,^8, 9^ this pathophysiological correlate seems to hold true for SID in patients with essential tremor.

### Methodological considerations and limitations

While previous studies focused on phonematic and articulographic features of SID^8, 9, 11, 13^ this study expands these findings by (1) revealing the structural network underlying SID and (2) providing an atlas structure predictive of stimulation-induced changes in intelligibility. Furthermore, this study highlights the spatial proximity of tremor control and SID, which is often experienced in clinical practice and may lead to suboptimal postoperative tremor control. Although the cohort-size of fourteen patients might seem small, a total of 111 stimulation settings could be included in the analysis. To estimate the VTAs based on these stimulation settings, a well-established approach was used, that has previously been employed to identify optimal target regions and connectivity profiles in Parkinson’s disease and essential tremor.^22, 25, 31, 32, 42^ However, the theoretical concept of VTA modelling constitutes a simplification that neglects factors like fiber orientation.^43^ Regarding the concept of predicting effects of deep brain stimulation in relation to the anatomical target area there have been several approaches in the past, such as investigating lead positions,^1^ the creation of probabilistic sweet spots,^31, 42^, and the estimation of beneficial connectivity profiles ^22, 32^. In this study, we focused on providing an atlas structure that is easy to implement in planning and programming software and therefore decided to base our predictive model on a well-established approach introduced by Baldermann et al..^32, 33, 44, 45^ In these studies including clinical stimulation settings only, T-tests and T-statistics were employed to define discriminative fibers and to predict the respective outcome parameter. In contrast, in the present study determination of discriminative fibers was based on linear mixed effect models to control for multiple testing per patient. Some methodological limitations should be considered when interpreting the results of this study. First, the sample size of the present study is limited to 14 patients, which is within the range of previous studies examining deterioration of the speech motor system in essential tremor patients with thalamic DBS.^8, 9, 13, 17^ In contrast to these studies mainly comparing “OFF stimulation” state to “ON stimulation” state, the present study included 10 test settings per patient. Additionally, despite the reduced spread of patients with varying severities of SID, the results held true in the leave-one out analysis. Second, VTAs were pooled on the left hemisphere for the analysis. This approach seems appropriate as all patients were right-handed and previous studies did not show any difference between right and left hemispheric DBS regarding SID in bilaterally implanted essential tremor patients.^11^ However, there are studies reporting SID to be associated with left-hemispheric stimulation of the subthalamic nucleus in Parkinson’s disease.^46, 47^ Third, an atlas-based approach neglects a certain degree of inter-individual heterogeneity, but the employed state-of-the-art multispectral co-registration approach has proved to be accurate.^26, 48^ Fourth, this study employed normative connectome data to estimate structural connectivity in individual patients. This approach has already been implemented in many studies investigating the effects of DBS when patient-specific imaging data allowing fiber tracking was lacking, e.g., in Parkinson’s disease and obsessive-compulsive disorder,^22, 33^ but also essential tremor.^32, 49^ While these connectomes do not represent patient-specific connectivity, they are superior in quality to most images acquired during clinical routine by using specialized magnetic resonance hardware and best-performing tractography processing algorithms.^50^ Most importantly, using this specific normative connectome data allowed us to compare our results regarding SID to the results by Al-Fatly et al.^32^ regarding tremor suppression without being biased by different imaging acquisition protocols. Lastly, interindividual anatomical variability and center-specific targeting strategies might lead to different target points. This raises the question of the generalizability of our results to other cohorts. Therefore, future studies, including different centers and target points, are warranted to create more generally applicable predictive models.

## Conclusion

The present study demonstrates that both, the motor cortex as well as the cerebellum are involved in stimulation-induced speech impairment in essential tremor patients. Furthermore, the derived fiber-based atlas structure might help to avoid stimulation-induced dysarthria in essential tremor patients and could easily be implemented in postoperative programming strategies.

## Abbreviations

DBS: deep brain stimulation
DDK: diadochokinesis
SID: stimulation-induced dysarthria
PSA: posterior subthalamic area
STN: subthalamic nucleus
TRS: tremor rating scale
VIM: ventral intermediate nucleus
VTA: volume of tissue activated

## Acknowledgments

The authors thank Bassam Al-Fatly and coworkers for providing the fibers associated with tremor control as published in Al-Fatly et al. 2019. In remembrance of Johannes Becker, whose enthusiasm, steadiness and criticism enriched this scientific field.

## Footnotes

### Author contributions

JNPS, HJ, TT, TAD, and MTB contributed to the conception and design of the study; JNPS, HJ, TT, JKS, PR, HSD, DM, GRF, VVV, TAD and MTB contributed to the acquisition and analysis of the data; JNPS, HJ and TAD, contributed to drafting the text and preparing the figures. All authors reviewed and revised the manuscript for intellectual content.

